# Efficient Bayesian inference for ordinary differential equation models from experimental data with uncertain measurement times

**DOI:** 10.64898/2026.05.09.724053

**Authors:** Jakob Vanhoefer, Vanessa Nakonecnij, Nadine Binder, Jan Hasenauer

## Abstract

Time-resolved measurements are central to calibrating mechanistic dynamical models, but current inference frameworks typically assume that reported measurement times are exact. In practice, actual sampling times may deviate from reported times because of sample-handling delays, imperfect synchronization, or reporting errors. Here, we present a Bayesian framework for parameter inference in ordinary differential equation models that explicitly accounts for uncertainty in measurement times. We formulate latent measurement times as random variables and derive a joint and marginalized posterior. To compute the marginal likelihood efficiently, we augment the original dynamical system with additional state variables that evaluate the required integrals during numerical simulation. This reduces the dimensionality of the estimation problems and allows for efficient and reliable Markov chain Monte Carlo sampling. Across synthetic examples and a published model of carotenoid cleavage in *Arabidopsis thaliana*, neglecting time uncertainty led to biased estimates and overconfident uncertainty quantification, whereas the proposed marginalized formulation recovered reliable parameter estimates while substantially improving sampling efficiency and scalability. These results identify measurement time uncertainty as an important source of variability in dynamic modeling and establish posterior marginalization as a practical strategy for robust mechanistic inference.

## 1 Introduction

Dynamical mathematical models provide a quantitative framework for describing and analyzing the temporal evolution of biological systems (Kitano, 2002). By formulating mechanistic hypotheses in terms of state variables and their interdependence, such models enable the integration of heterogeneous experimental data, the systematic investigation of causal relationships, and the prediction of system responses under novel conditions. Among the different classes of dynamical models, ordinary differential equation (ODE) models constitute the most widely used approach for describing cellular and biochemical processes (Klipp et al., 2005; Fall et al., 2010; Hasenauer et al., 2015). ODE models describe the time evolution of the concentrations of biochemical species based on reaction rates and stoichiometry, thereby establishing an explicit link between biochemical mechanisms and observable quantities. This mechanistic structure facilitates the quantitative testing and falsification of competing hypotheses, the characterization of process dynamics, and the rational design of experiments and perturbations (Faller et al., 2003; van Riel, 2006; Fröhlich et al., 2019). Consequently, ODE models provide a principled and widely adopted framework for the analysis of time-resolved measurements in systems and computational biology. Yet, ODE models typically contain unknown parameters, including reaction rate constants, transport rates, or initial conditions.

Some model parameters might be available in the literature or in public repositories such as BRENDA (Chang et al., 2009) or SABIO-RK (Wittig et al., 2018). Yet, the remaining parameters need to be inferred from experimental data. A broad spectrum of statistical methods for parameter estimation is available, including frequentist approaches based on maximum likelihood estimation and Bayesian approaches based on posterior inference; see the work by Raue et al. (2013) and references therein. These methods are implemented in established toolboxes such as Data2Dynamics (Raue et al., 2015) and pyPESTO (Schälte et al., 2023), and calibration problems can be formulated in standardized formats using Systems Biology Markup Language (SBML) (Hucka et al., 2019) and PEtab (Schmiester et al., 2021). Existing methodologies support a wide range of measurement noise models, including additive and multiplicative noise formulations, and provide strategies for handling outliers to ensure robustness (Loos et al., 2018).

Despite this methodological maturity, current approaches predominantly account for uncertainty in the measured quantities (Balsa-Canto et al., 2025), while treating the measurement time points as known and fixed. This assumption is suboptimal in many experimental settings. For instance, sampling procedures may require a non-negligible duration, quenching or blocking solutions may be applied with delays, or measurements may reflect processes that are not synchronized across experimental units. As a consequence, reported measurement times can be subject to substantial uncertainty, which is not explicitly represented in standard inference frameworks.

A straightforward strategy for handling uncertain measurement times is to treat the latent measurement time points themselves as optimization variables. However, this increases the dimensionality of the inference problem and can substantially worsen numerical performance. In related settings, namely for scaling and offset parameters, such dimensionality inflation has been addressed using hierarchical optimization. For example, Loos et al. (2018) and Schmiester et al. (2019) introduced hierarchical optimization approaches in which analytically tractable inner problems reduce the effective dimension of the outer optimization. However, these approaches benefit substantially from the analytical tractability of the inner optimization problems, which is not available in the present setting.

A related but more general strategy is marginalization, that is, integrating out nuisance variables instead of estimating them jointly. This approach is well established in survival analysis for interval-censored data and dates back to the foundational work of Turnbull (1976) and Sun (2006). However, these methods typically assume relatively simple parametric or semi-parametric event time distributions and do not scale to the high-dimensional integration problems arising in ODE models with many uncertain observation times and nonlinear dynamics. In the context of dynamical systems, Raimúndez et al. (2023) proposed a framework that marginalizes over scaling and offset parameters and derived closed-form expressions for the resulting marginalized likelihood. However, such closed-form expressions for the respective integrals are not known for uncertain measurement time points and are unlikely to exist for arbitrary temporal dynamics.

In this study, we present a Bayesian framework for modeling uncertainty in measurement times (Figure 1). Based on this formulation, we derive an efficient marginalization approach that removes the temporal parameters from the sampling problem while preserving their contribution to the posterior distribution. The proposed method augments the dynamical system such that the required marginal integrals can be evaluated alongside the state trajectory, enabling efficient and error-controlled numerical integration as well as sensitivity analysis using forward and adjoint methods. We assess the approach using synthetic examples and an application to experimental data. The marginalized formulation substantially reduces the effective dimensionality of the inference problem and improves convergence and sampling efficiency compared with joint estimation of dynamic and temporal parameters.

**Figure 1:**
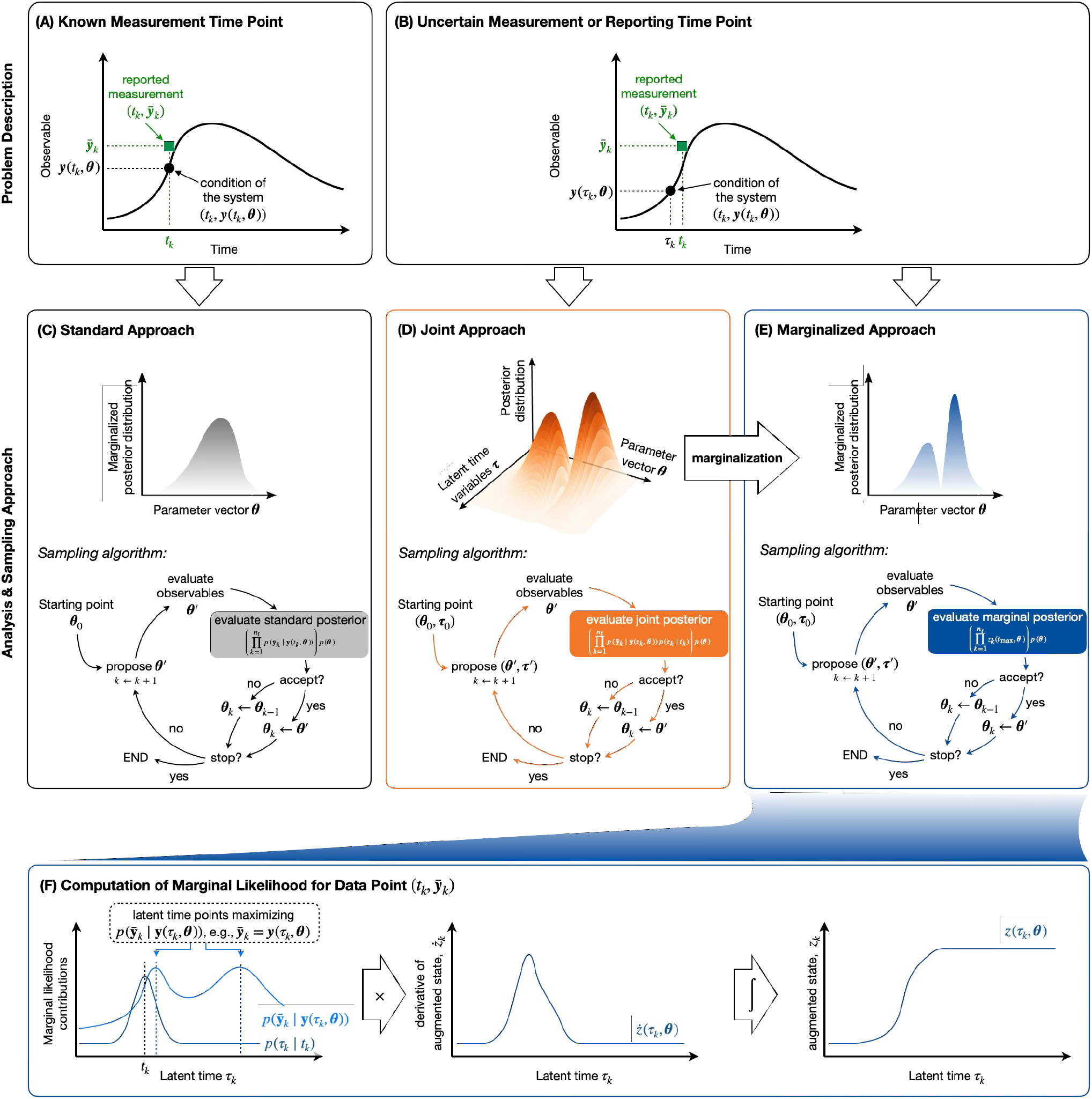
Measurement setups and analysis approaches. (A,B) Illustration of measurement settings with exactly known time points and with uncertain measurement or reporting times, showing the relationship between reported times *t*_*k*_, latent times *τ*_*k*_, model predictions ***y***(*t*_*k*_, ***θ***) and ***y***(*τ*_*k*_, ***θ***), and observations 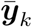. (C–E) Overview of the standard approach for known measurement times, and the joint as well as the proposed marginalized approach for uncertain measurement or reporting times, including corresponding Markov chain Monte Carlo schemes. (F) Computation of the marginal likelihood contribution for a single observation by integrating the product of the time prior and the observation likelihood, using an augmented state variable *z*_*k*_ that accumulates the integral during ODE simulation.

## 2 Methods

### 2.1 Mathematical modeling of biological processes

We consider ordinary differential equation (ODE) models of biochemical reaction networks, which are widely used to describe cellular processes such as signal transduction, gene regulation, and metabolism. The concentrations of chemical species are denoted by 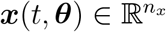 and satisfy

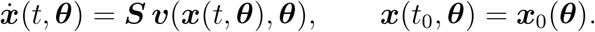

Here, 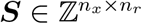 denotes the stoichiometric matrix and 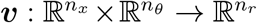 the vector of reaction fluxes. The parameter vector 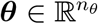 contains kinetic parameters such as reaction rate constants and may additionally include parameters determining the initial condition ***x***_0_(***θ***).

### 2.2 Mathematical modeling of the measurement process

In practice, only partial observations of the state vector ***x***(*t*, ***θ***) are available. Measurements may correspond to individual state variables, linear combinations of states (e.g., aggregated concentrations obtained from antibody-based assays), or nonlinear functions of the state (e.g., due to saturation effects). We therefore introduce an observable function 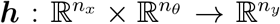, which maps the state vector to the observable vector according to

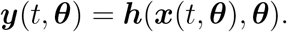

For each measurement, a measurement time point *t*_*k*_ and a measured value 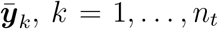, is reported. The measurements are assumed to be corrupted by noise. The dataset is given by

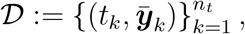

with *n*_*t*_ denoting the total number of measurements.

In the following, we distinguish two types of measurement-time assumptions.

#### Exactly known measurement time points

In the classical setting, measurements are assumed to be collected exactly at the reported time points *t*_*k*_ (Figure 1A). Assuming independence between measurements at different time points, the measured values follow

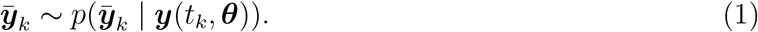

For independent, normally distributed measurement noise, this corresponds to

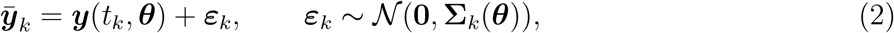

where **Σ**_*k*_(***θ***) denotes the noise covariance matrix. If measurement noise is independent across observables, the covariance simplifies to 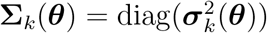.

Under the assumption of independence across time points, the likelihood function becomes

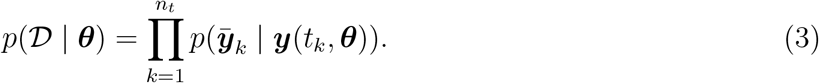

In the Gaussian noise case with independence across time points, this reduces to the product of multivariate normal densities,

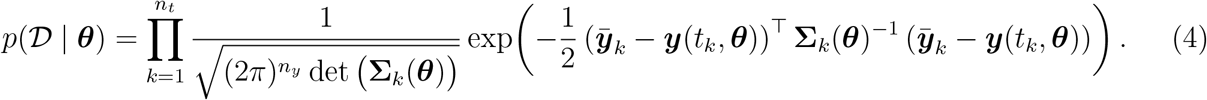

#### Uncertain measurement or reporting time points

In addition to the classical setting, we consider the case where measurement or reporting time points are subject to uncertainty (Figure 1B). Two conceptually different scenarios can be distinguished:

i. The experiment is planned to be conducted at a time *t*_*k*_, yet, due to variability in the experimental process (e.g., sample handling), the actual measurement time *τ*_*k*_ may deviate from *t*_*k*_. In this case, the measurement time is a random variable which is conditioned on the planned/reported time *t*_*k*_,

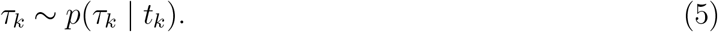

This scenario is typical for well-controlled experimental settings, such as basic science experiments.
ii. The experiment is conducted at a time point *τ*_*k*_, but the reported time point *t*_*k*_ is subject to inaccuracies in the recording process. In this case, the reported time *t*_*k*_ is modeled as a random variable which is conditioned on the measurement time *τ*_*k*_,

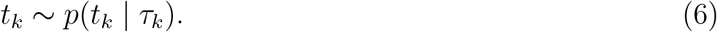

This scenario is more common in applied settings, such as clinical studies, where the reporting process may be partially decoupled from the actual data acquisition.

These two scenarios differ only in the direction of the conditional distribution but describe the same underlying relationship between actual and reported time points. In both cases, a probability distribution characterizes the discrepancy between measurement and reporting times. In the following, we adopt the first formulation, *p*(*τ*_*k*_ | *t*_*k*_), for notational convenience; however, all results can be equivalently formulated under the second perspective.

Given the latent measurement time *τ*_*k*_, the reported measurement 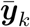 corresponds to the system state at time *τ*_*k*_:

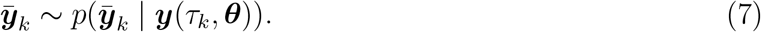

Conditioned on the latent time variables 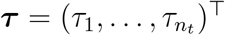, the joint likelihood factorizes as

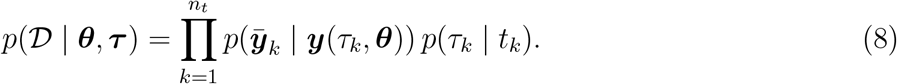

### 2.3 Parameter estimation and uncertainty quantification

The mathematical models describing the biological and measurement processes contain unknown parameters ***θ***, including reaction rate constants, initial conditions, and measurement-specific parameters such as scaling, offset, and noise terms. We estimate these parameters within a Bayesian framework based on Bayes’ theorem.

#### Exactly known measurement time points

In the classical setting with exactly known measurement time points, the posterior distribution of the parameters is proportional to the product of likelihood *p*(𝒟 | ***θ***) and prior distribution *p*(***θ***),

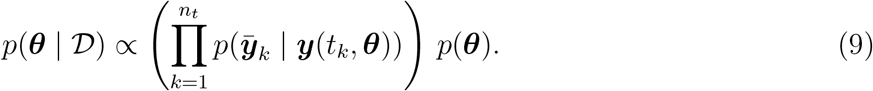

Point estimates are obtained via maximum a posteriori (MAP) estimation. Since the marginal likelihood does not depend on ***θ***, the MAP estimator is obtained by minimizing

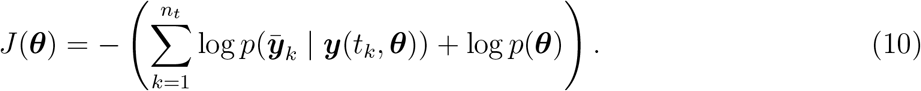

which is the negative log-posterior up to an additive constant. The resulting optimization problem

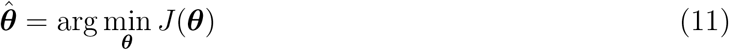

is generally nonlinear and non-convex. To address this, we employ multi-start local optimization. Parameter uncertainty is quantified by sampling from the posterior distribution using Markov chain Monte Carlo (MCMC) methods.

In the following, we refer to optimization and uncertainty quantification using the standard likelihood without unknown measurement or reporting time points as the **standard approach** (Figure 1C).

#### Uncertain measurement time points

When measurement time points are subject to uncertainty, the actual sampling times 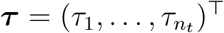 are treated as additional unknown variables. The joint posterior distribution of parameters and measurement times is proportional to the product of likelihood and prior distribution,

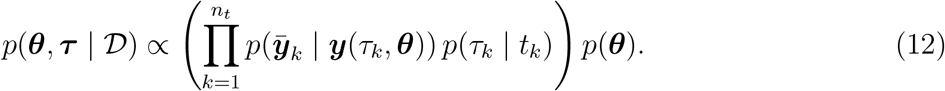

A MAP estimate of parameters and time points is obtained by minimizing

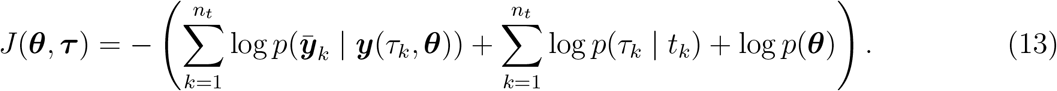

The corresponding optimization problem,

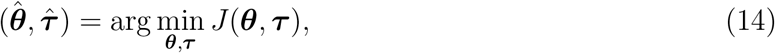

is—as before—generally nonlinear and non-convex. In addition, the dimension of the optimization problem, *n*_*θ*_ +*n*_*t*_, grows with the number of observations. For dense time series, *n*_*t*_ can substantially exceed *n*_*θ*_. While gradient-based local solvers often require a number of iterations that is only weakly dependent on the number of optimization variables (Hass et al., 2019), the computational cost per iteration and the complexity of the optimization landscape can still increase markedly.

Similar issues arise for posterior sampling. Although adaptive MCMC methods can learn aspects of the posterior geometry, high-dimensional settings with pronounced correlation structures between ***θ*** and ***τ*** may exhibit slow mixing and reduced effective sample size.

In the following, we refer to optimization and uncertainty quantification based on the joint posterior distribution as the **joint approach** (Figure 1D).

### 2.4 Marginalization over latent measurement time points

To mitigate the challenges associated with the optimization and sampling of the potentially high-dimensional joint posterior *p*(***θ, τ*** | 𝒟), we propose to marginalize over the latent measurement times ***τ***. This strategy is conceptually related to previous work on posterior marginalization of scaling, offset, and noise parameters for dynamical models (Raimúndez et al., 2023), where dimensionality reduction led to substantial gains in sampling efficiency. Here, we integrate out ***τ*** to obtain the marginalized posterior *p*(***θ*** | 𝒟) over ***θ***.

We assume that the distribution of the actual measurement time has compact support [*t*_0_, *t*_max_], i.e.,

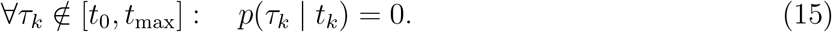

This assumption is obviously satisfied in practice.

#### 2.4.1 Marginalized posterior

Marginalizing the posterior with respect to the latent time variables ***τ*** over the interval [*t*_0_, *t*_max_] yields the marginalized posterior,

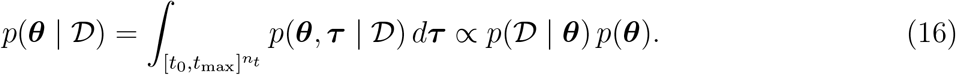

The marginal likelihood *p*(𝒟 | ***θ***) follows from (12) as

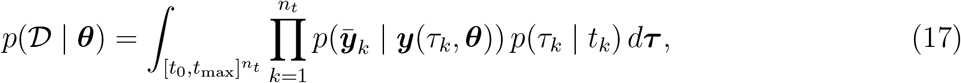

which is a high-dimensional integral and therefore difficult to evaluate directly. Fortunately, multiplication and integration can be interchanged, yielding a product of one-dimensional integrals,

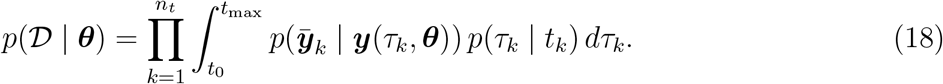

The individual one-dimensional integrals correspond to the contributions of different measurement time points to the marginal likelihood. The contribution of a measurement reported at time *t*_*k*_ is obtained by marginalizing the likelihood 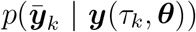 over the latent measurement time *τ*_*k*_, thereby accounting for the distribution *p*(*τ*_*k*_ | *t*_*k*_).

The individual integrals, as well as their product, can usually not be computed in closed form. Indeed, the integrand 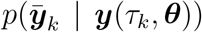 can have a complex structure. Multimodality might, for instance, occur if there exist multiple time points *τ*_*k*_ for which 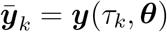.

#### 2.4.2 Computation of marginal likelihood

The individual contributions to the marginal likelihood can be computed by direct numerical quadrature. For this purpose, the trajectory ***y***(*t*, ***θ***) is stored during the forward solution of the ODE model. Using these stored values, quadrature schemes with predetermined quadrature points (e.g., the trapezoidal rule) can be applied. This approach is used as a baseline in the following. However, it does not allow for direct control of the integration error, and the resulting conservative step-size selection may lead to computational inefficiencies.

To ensure reliable and efficient computation of the contributions to the marginal likelihood, we instead propose to augment the original ODE model. Specifically, we introduce additional state variables *z*_*k*_(*t*, ***θ***), *k* = 1, …, *n*_*t*_, each defined via a one-dimensional ODE,

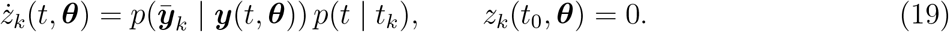

The value of these state variables at time *t*_max_ equals the marginalized likelihood contributions,

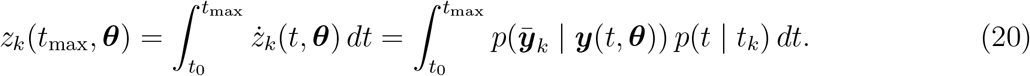

The state variables 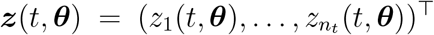 are computed alongside the state of the biological process ***x***(*t*, ***θ***) by solving the augmented ODE system

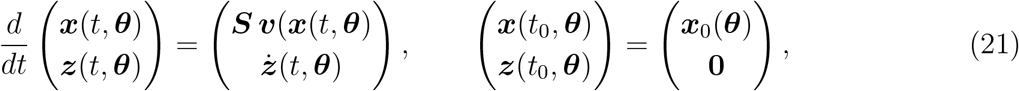

where 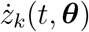 is defined according to (19). Solving the augmented system up to *t*_max_ yields ***z***(*t*_max_, ***θ***), from which the marginalized likelihood can be computed as

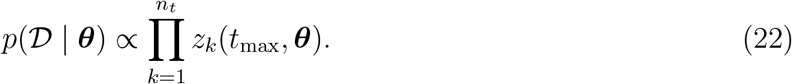

#### 2.4.3 Parameter estimation and uncertainty quantification

The marginalized posterior distribution enables both optimization and uncertainty quantification. In particular, the formulation using the augmented ODE model allows for the application of established forward and adjoint methods for gradient computation.

A marginal maximum a posteriori (MAP) estimate is obtained by minimizing the negative log-marginal posterior,

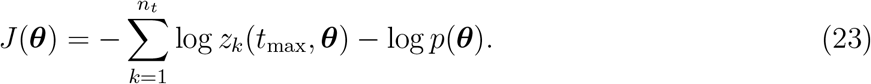

The corresponding optimization problem,

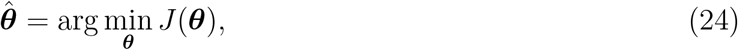

has dimensionality *n*_*θ*_ and is therefore lower-dimensional than the joint estimation problem (14), but generally remains nonlinear and non-convex. We solve it using the same multi-start local optimization strategy as described above. Importantly, since the latent measurement time points are integrated out—rather than maximized over—the marginal MAP estimate for ***θ*** obtained by solving (24) may differ from the joint MAP estimate obtained from (14).

Bayesian uncertainty estimates for ***θ*** are obtained by sampling from the marginalized posterior distribution using MCMC methods, employing the sampling algorithms and convergence diagnostics outlined above. Since the marginalization is exact up to tightly controlled numerical integration error, sampling from the marginalized posterior should, in principle, yield the same estimates as sampling from the joint posterior followed by discarding the latent measurement times. However, as sampler characteristics (e.g., proposal scaling) as well as general performance depend on dimensionality, the finite-sample behavior may—as demonstrated by Raimúndez et al. (2023)—differ.

Overall, marginalization over ***τ*** avoids explicit sampling of the latent variables and reduces the effective dimensionality of the inference problem. In analogy to the marginalization of measurement-specific parameters (Raimúndez et al., 2023), this reduction can substantially improve optimization and sampling performance, particularly when ***θ*** and ***τ*** are strongly coupled or when the posterior distribution is multimodal.

In the following, we refer to optimization and uncertainty quantification based on the marginalized posterior distribution as the **marginalized approach** (Figure 1E). The marginal likelihood calculation is illustrated in Figure 1F.

### 2.5 Implementation

Mathematical models of the biological processes were specified in the SBML (Keating et al., 2020) using yaml2sbml (Vanhoefer et al., 2021). Numerical simulation and sensitivity analysis were performed using the Advanced Multi-language Interface for CVODES and IDAS (AMICI) (Fröhlich et al., 2021), which builds on the SUNDIALS solvers CVODES and IDAS (Serban and Hindmarsh, 2005).

Objective functions were implemented as custom objectives in the Python Parameter Estimation Toolbox (pyPESTO) (Schälte et al., 2023). Parameter optimization was carried out using the trustregion method `fides` (Fröhlich and Sorger, 2022) via the pyPESTO interface. Posterior sampling was performed using the pyPESTO implementation of the adaptive Metropolis–Hastings algorithm (Haario et al., 2001). Burn-ins are removed using the iterative scheme outlined in (Ballnus et al., 2017) and implemented in pyPESTO. Convergence is assessed using the Geweke diagnostic and the Gelman–Rubin statistic.

The code used in this study is available on Zenodo under the BSD 2-Clause License.

## 3 Results

We illustrate the proposed approach for handling measurement time uncertainties using two in silico examples and one published model. Across the case studies, we compare three inference strategies introduced above: (i) a standard approach assuming that measurement time points are reported without error, (ii) a joint estimation approach that infers both model parameters and latent measurement times, and (iii) the proposed marginalized approach that infers model parameters by integrating out latent measurement times using the augmented ODE formulation.

### 3.1 Neglecting temporal uncertainty leads to biased and overconfident estimates

To illustrate the impact of measurement time uncertainty on parameter inference and how different formulations handle it, we consider a minimal exponential decay model,

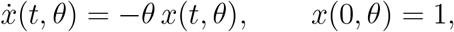

with decay rate *θ* and observable *y*(*t, θ*) = *x*(*t, θ*). We assumed a uniform prior, *p*(*θ*) ∼ 𝒰(0, 5). Synthetic data were generated using *θ*^true^ = 1.0 and additive Gaussian noise with standard deviation *σ*_*y*_ = 0.1. Measurements were reported at time points *t* = [0.5, 2.0], while the true measurement times were sampled using a truncated normal distribution,

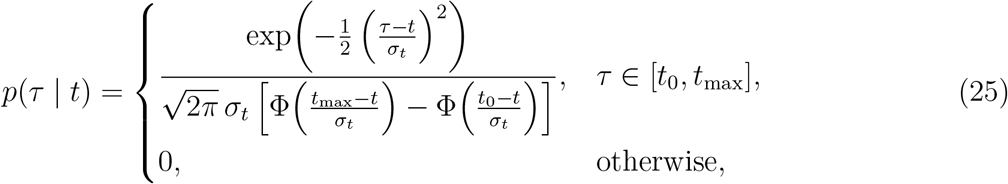

where Φ denotes the cumulative distribution function of the standard normal distribution, with *t*_0_ = 0, *t*_max_ = 10, and *σ*_*t*_ = 0.5. For demonstration, we intentionally focus on a low-sample regime with two observations and substantial temporal uncertainty (Figure 2A).

**Figure 2:**
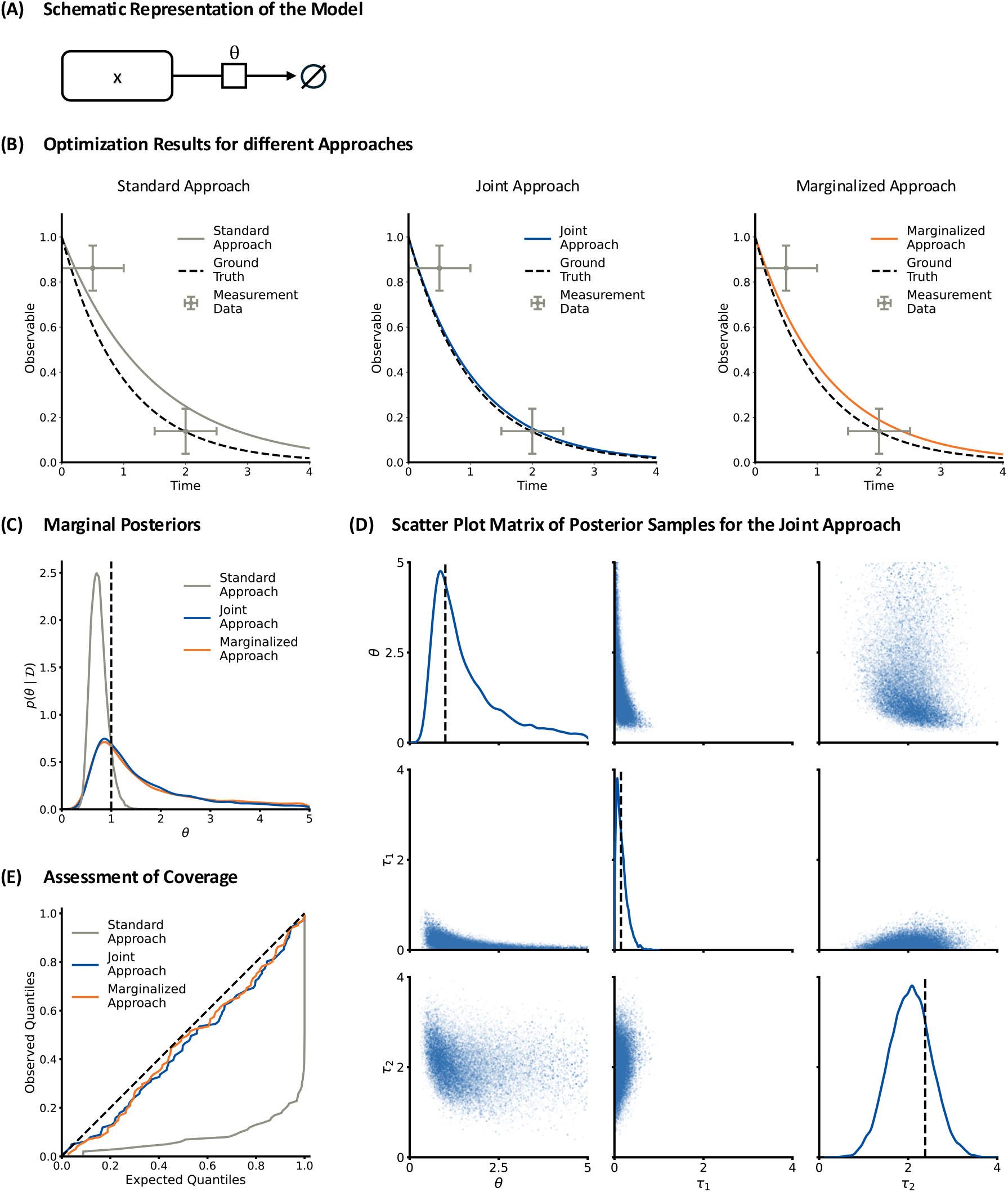
Illustration of the impact of measurement time uncertainty using an exponential decay model. (A) SBGN diagram of the exponential decay model. (B) Synthetic data and model fits obtained using the standard, joint, and marginalized approach. (C) Marginal posterior distributions of the decay rate *θ* obtained from the standard, joint, and marginalized approaches. (D) Scatter plot of posterior samples for the joint formulation. (E) Probability-probability plot for theoretical vs. empirical cumulative distribution function (CDF). The cumulative density functions correspond to the observed coverage for 100 synthetic datasets. (A–D) Results for the same synthetic dataset, and (E) is based on 100 synthetic datasets.

We evaluated the standard, joint, and marginalized formulations of the estimation problem. MAP estimates and sampling results show that neglecting temporal uncertainty leads to inaccurate estimates of the decay dynamics (Figure 2B) and the decay rate (Figure 2C), while both joint and marginalized approaches provide more accurate estimates. Indeed, the marginal posterior distributions obtained from the joint and marginalized approaches closely align (Figure 2C), confirming the consistency of the two formulations. Furthermore, sampling of the joint posterior reveals a pronounced dependency between *θ* and the latent time variables (Figure 2D), indicating a nontrivial posterior geometry.

To assess the reliability of the uncertainty quantification, we repeated data generation and sampling for 100 synthetic datasets. For each run, we computed credible intervals and evaluated their coverage of *θ*^true^. The resulting probability-probability plot (Gibbons and Chakraborti, 2014) shows that ignoring temporal uncertainty leads to severe overconfidence, with substantially underestimated credible intervals (Figure 2E). Indeed, the 80% credible interval computed using the standard approach contains *θ*^true^ only for 13 of 100 synthetic datasets, compared to 73 and 75 synthetic datasets for the joint and the marginalized approaches.

These results show that for reliable and predictive dynamical modeling a thorough handling of time uncertainties is critical, and that the joint and the marginalized formulation provide a good starting point.

### 3.2 Posterior marginalization enables efficient sampling

To investigate the computational implications of the different inference strategies, we consider a two-state model (Figure 3A),

**Figure 3:**
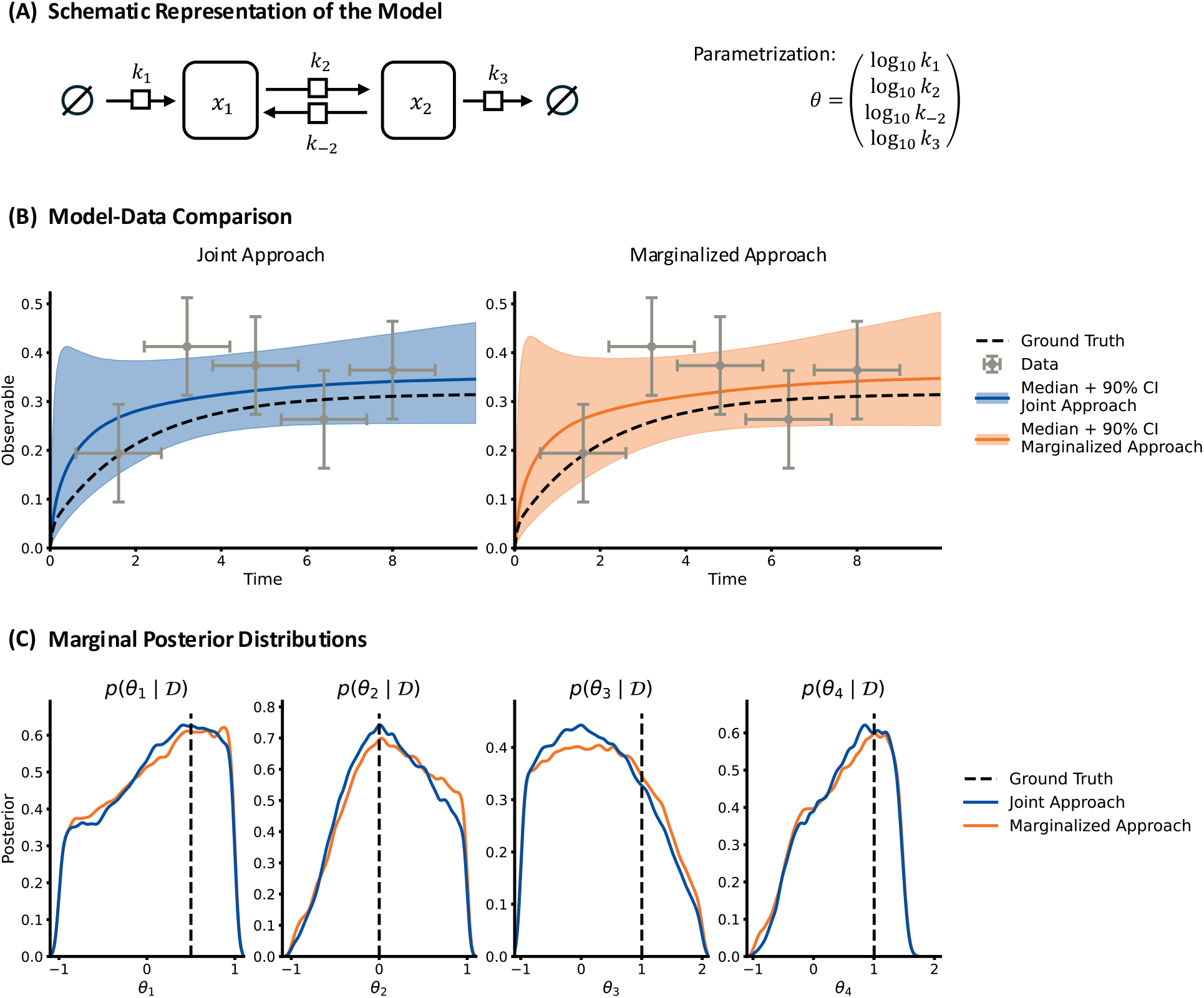
Comparison of sampling results for 2-state model. (A) SBGN diagram of the two-state model. (B) Comparison of synthetic data and model predictions obtained from posterior samples for the joint and marginalized approaches. (C) Marginal posterior distributions of the model parameters for the joint and marginalized formulations. (A–C) Results are shown for the same synthetic dataset with *n*_*t*_ = 4 time points.

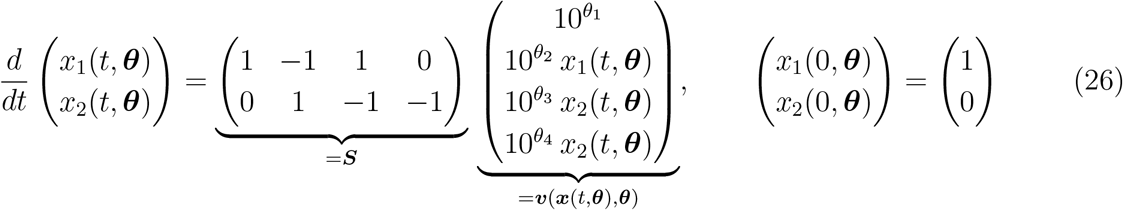

with synthesis rate *θ*_1_, conversion rates *θ*_2_ and *θ*_3_, and a degradation rate *θ*_4_. Synthetic data for the observable *y*(*t*, ***θ***) = *x*_2_(*t*, ***θ***) were generated using ***θ***^true^ = (0.5, 0, 1, 1)^⊤^. Measurement noise was assumed to be Gaussian with standard deviation *σ*_*y*_ = 0.1. Measurements were reported at equidistant time points, while the true measurement times were sampled from a truncated normal distribution with *t*_0_ = 0, *t*_max_ = 12, and *σ*_*t*_ = 1. We assumed a log-uniform prior, *p*(***θ***) ∼ 𝒰_log_(−1, 2), and performed MCMC sampling with 5 × 10^5^ iterations per chain.

For a small number of observation time points (e.g., *n*_*t*_ = 4), both the joint and marginalized approaches yield similar posterior distributions and accurately describe the data (Figure 3B). The marginal posterior distributions of the mechanistic parameters are in close agreement, indicating that both formulations recover consistent parameter estimates (Figure 3C).

Considering a broader range of observation time points reveals clear differences in computational performance. For the joint formulation, the number of unknown parameters increases linearly with the number of observations (*n*_*θ*_ + *n*_*t*_), whereas for the marginalized formulation the number of parameters remains constant (*n*_*θ*_), and only the number of augmented state variables increases (Figure 4A). This difference leads to markedly different MCMC behavior (Figure 4B-D): for the joint formulation, sampling performance deteriorates as the number of time points increases, whereas it remains largely unchanged for the marginalized formulation.

**Figure 4:**
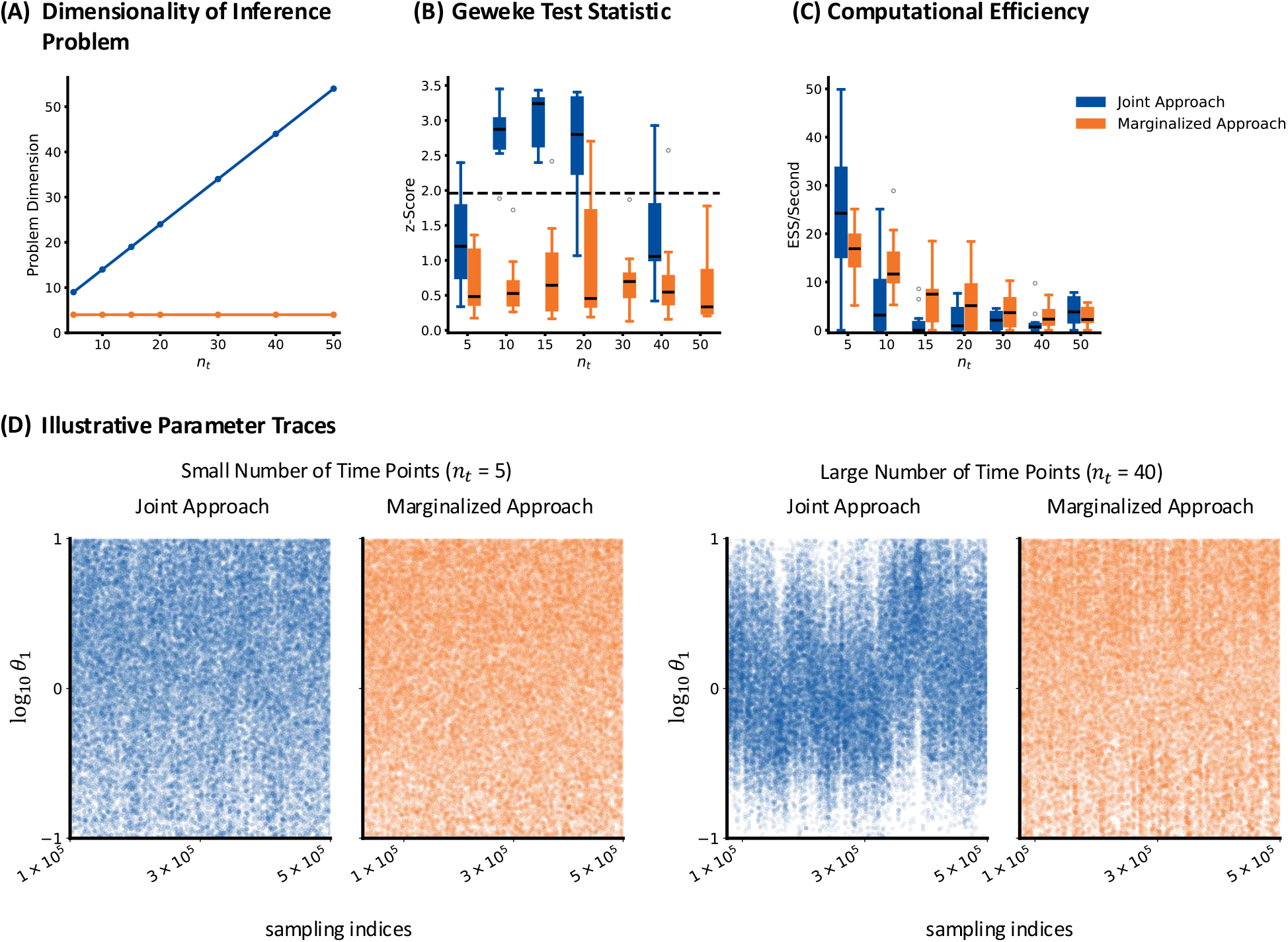
Assessment of scalability and sampling performance for 2-state model. (A) Dimensionality of the inference problem as a function of the number of observation time points for the joint and marginalized formulations. (B) Distribution of the z values of the Geweke test statistic for joint and marginalized approach, and threshold which would be considered significant for rejecting a run. (D) Effective sample size per unit computation time as a function of the number of observation time points. (C) Representative MCMC traces for small and large numbers of observation time points, illustrating the deterioration of sampling performance for the joint formulation. (B–C) Results based on 10 synthetic datasets pewith varying numbers of observations, and (D) a single synthetic datasets.

The analysis of the MCMC chains using established convergence diagnostics, such as Geweke statistics, reveals that many chains for the joint formulation fail to converge (Figure 4B). Accordingly, the effective sample size per unit computation time drops for the joint approach, while it remains relatively stable for the marginalized approach (Figure 4C). These quantitative assessments are consistent with visual inspection of the MCMC chains (Figure 4D), and show that the posterior marginalization substantially improves sampling efficiency and scalability.

### 3.3 Sampling inaccuracies can be detected in a real-world model

To assess the applicability of the proposed approach, we consider a published model of carotenoid cleavage in *Arabidopsis thaliana* (Bruno et al., 2016) (Figure 5A). The model describes enzymatic cleavage reactions of carotenoids and has been widely used for benchmarking dynamic modeling approaches. It comprises seven state variables and 13 parameters, including kinetic rates and condition-specific initial values. Multiple observables corresponding to measured metabolite concentrations are available across several experimental conditions. Further details on the model structure, observables, and experimental setup are provided in the original publication. For our analysis, we adopt the same parameter bounds as provided in the available PEtab implementation (Schmiester et al., 2021) and use a log-uniform parameter prior.

**Figure 5:**
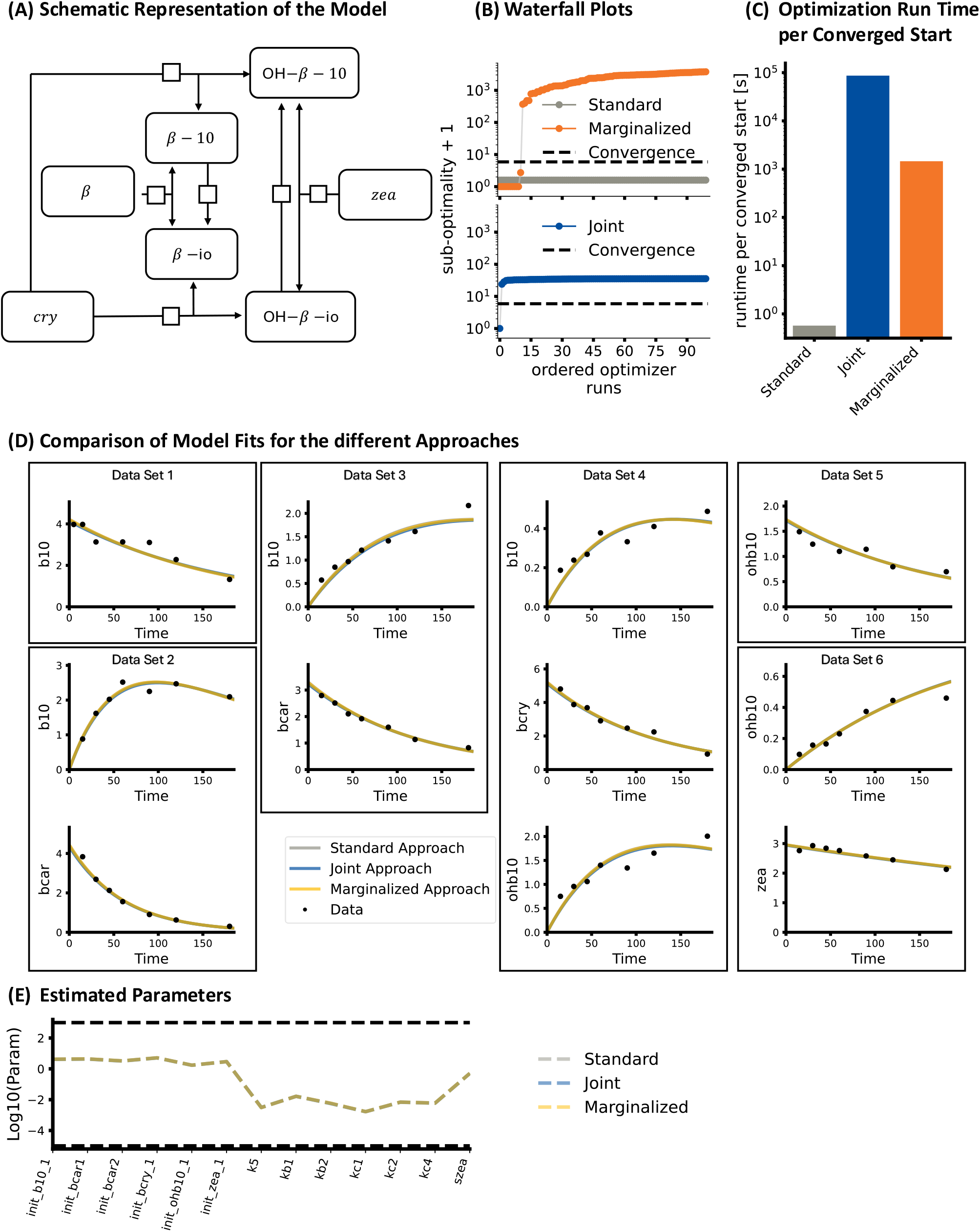
Parameter estimation for a published carotenoid cleavage model and dataset. (A) Schematic representation of the carotenoid cleavage pathway in *Arabidopsis thaliana*. (B) Waterfall plots of sorted objective function values from multi-start optimization using the standard, joint, and marginalized approaches. The value on the y-axis indicates the suboptimality, which is the difference from the best-found objective function value, plus one. As the objective function of the marginalized approach simplifies to the objective function of the standard approach for *σ*_*t*_ = 0, they are plotted together, with the best objective function values found using the marginalized approach providing the optimum. (C) Computation time per converged start (as defined by *J* − *J* ^opt^ *<* CDF*χ*^2^(0.95)) for the standard, joint, and marginalized approaches. (D) Comparison of measurement data and model predictions obtained from the MAP estimates for the standard, joint, and marginalized approaches. (E) Parallel coordinates plot of MAP estimates for the standard, joint, and marginalized approaches. (B-E) Results are shown for the original dataset without added reporting time errors; nevertheless, the joint and marginalized approaches still estimate the reporting time error. Observable and parameter naming follows the PEtab implementation of the model (Schmiester et al., 2021).

The original dataset does not report measurement time uncertainties. Therefore, we treat the published measurement time points as a reference for the actual measurement time points and introduce artificial errors in the reported times. Specifically, we sample perturbed reporting times from truncated normal distributions with different standard deviations *σ*_*t*_ to emulate different levels of reporting error. In contrast to the previous examples, the scale parameter of the time uncertainty, *σ*_*t*_, is treated as unknown and estimated jointly with the remaining model parameters.

Our analysis of the model confirmed the finding of the previous section that parameter estimation using the joint approach is computationally demanding compared to the standard and marginalized approaches (Figure 5B & C). For the original data (*σ*_*t*_ = 0), the convergence rate of the joint approach was much lower than that of the standard approach (Figure 5B). Indeed, the joint approach found the respective best objective function value only once, while the marginalized approach provided several replicates, yielding a substantially better computation time per converged optimization (joint approach: 86703.68 seconds per converged start; marginalized approach: 1457.78 seconds per converged start; see Figure 5C). Yet, the marginalized approach was still computationally much more demanding than the standard approach due to: (1) a higher computational cost of the objective function and gradient evaluation, which is related to the dimensionality difference between the model (7 state variables) and the augmented model (7 + 77 state variables); and (2) a lower convergence rate per start. The fits for all methods agreed well with the experimental data (Figure 5D), and the parameter estimates were similar (Figure 5E). Indeed, the joint and marginalized approaches provided low estimates for *σ*_*t*_, which is expected as no noise was added. Due to the comparatively high computational cost and the similarity of the results, we consider only the standard and marginalized approaches in the following.

The analysis of the datasets with different reporting time errors revealed that the standard approach provides estimates that deviate substantially from the reference solution obtained using the unperturbed data (Figure 6A). While the estimation error also increased for the marginalized approach, the estimates remained sightly more consistent with the reference (Figure 6A). Indeed, the MAP estimate of the inferred reporting error increased with the applied reporting error level (Figure 6B). The computation of the likelihood ratio between the standard and the marginalized approach shows that larger reporting error levels can be clearly pinpointed (Figure 6C). Indeed, the analysis of the real data (*σ*_*t*_ = 0) indicates the absence of reporting time errors—as we expected— while for large perturbations(*σ*_*t*_ ≥ 5), the standard model is mostly rejected (Figure 6D).

**Figure 6:**
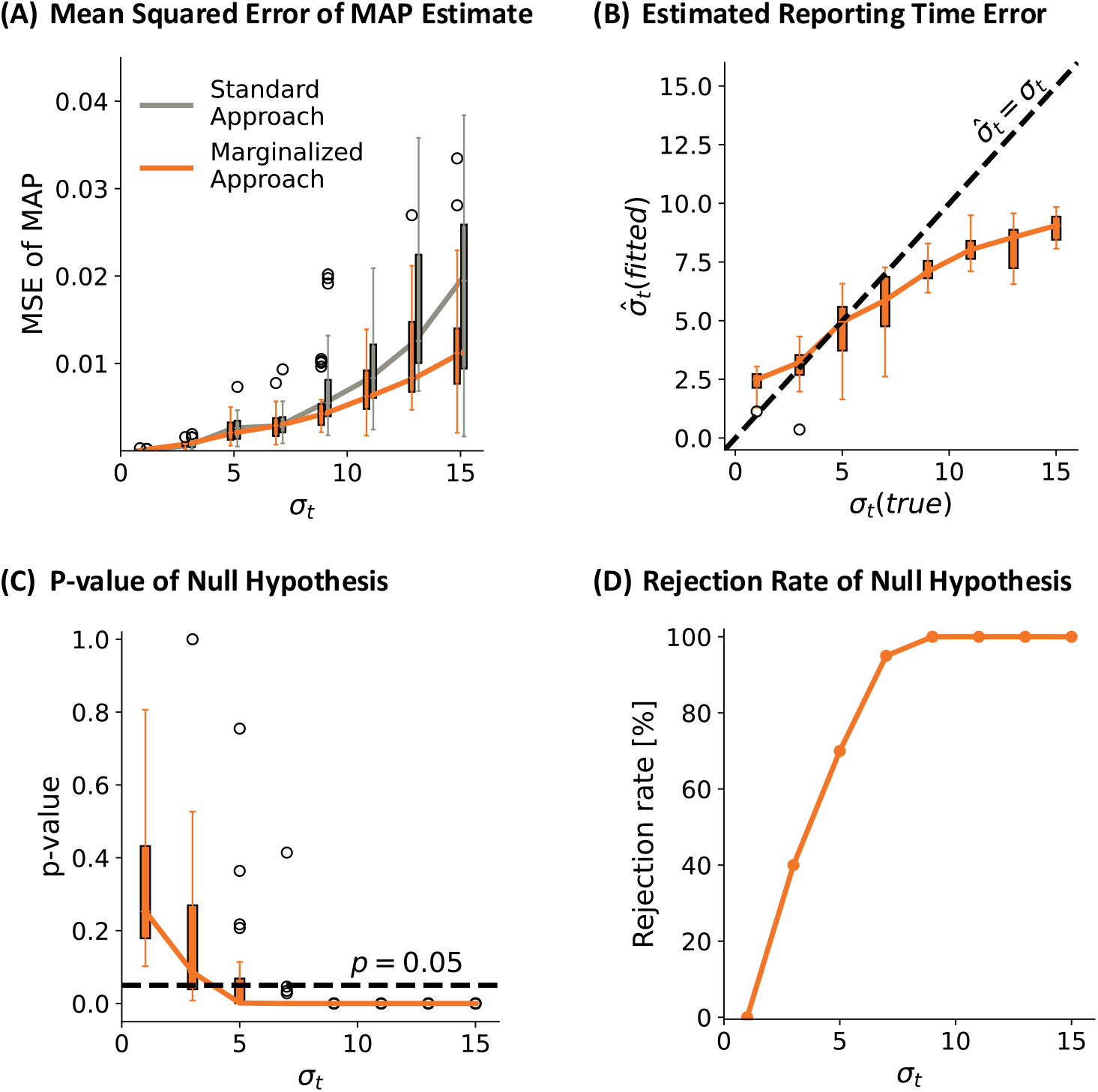
Detection and correction of measurement time uncertainty for a published carotenoid cleavage model and perturbed dataset. (A) Illustration of the impact of reporting time uncertainties on the MAP estimates for the standard and marginalized approaches. The Mean Squared Error (MSE) is computed between MAP estimate for the unperturbed data and the perturbed data, using the log_10_-transformed parameters. (B) Box plot of MAP estimates for reporting error levels obtained using marginalized approach. (C) Box plot of p-values of likelihood ratio test with the standard model as null hypothesis and the marginalized models as alternative hypothesis. The dashed line indicates rejection of the null hypothesis that there is no reporting error at an *α* level of 0.05. (D) Fraction of rejections of the null hypothesis at an *α* level of 0.05. (A-D) Results are computed / shown for 20 synthetic datasets.

These results demonstrate that the proposed approach is applicable to realistic models and enables the detection of and compensation for measurement time uncertainties in experimental data.

## 4 Discussion

In this work, we investigated the impact of measurement time uncertainty on parameter estimation and uncertainty quantification in dynamic models. We introduced a marginalized formulation that integrates out latent measurement times and compared it to a standard approach neglecting temporal uncertainty and a joint formulation estimating both parameters and latent time points.

Our results demonstrate that neglecting measurement time uncertainty can lead to biased parameter estimates and substantial underestimation of uncertainty. Even in simple models, this effect is pronounced and results in overconfident posterior distributions. In contrast, both the joint and marginalized formulations provide consistent parameter estimates and appropriate uncertainty quantification. These findings highlight that temporal uncertainty represents an important and often overlooked source of variability that should be accounted for in model-based analyses.

While the joint formulation provides a conceptually straightforward extension by treating measurement times as additional parameters, it leads to a substantial increase in dimensionality. As shown in our analyses, this increase adversely affects sampling performance, resulting in poor mixing and lack of convergence for larger datasets. In contrast, the marginalized formulation avoids this issue by keeping the parameter space fixed and incorporating temporal uncertainty through integration, leading to improved scalability and sampling efficiency while preserving statistical consistency with the joint formulation. The idea of marginalization builds on our previous work on nuisance parameters such as scaling, offset, and noise (Raimúndez et al., 2023). However, handling temporal uncertainty is substantially more challenging due to its nonlinear effect on the system dynamics.

The proposed formulation based on an augmented ODE system provides an elegant solution, as it enables the use of established simulation and inference toolboxes. Moreover, explicitly modeling time uncertainty may improve robustness to inaccuracies in reported measurement times, including mislabeled observations, by capturing the underlying mechanism rather than relying on heavy-tailed noise models (Maier et al., 2017).

The application to a published carotenoid cleavage model demonstrates that the proposed approach is suitable beyond synthetic examples. By introducing controlled perturbations of measurement times, we showed that accounting for temporal uncertainty improves the robustness of parameter estimation and yields uncertainty estimates that are consistent with reference solutions. Moreover, the inferred distributions of temporal noise parameters reflect the magnitude of the introduced perturbations, suggesting that the presence of measurement time uncertainty can, in principle, be detected from sufficiently informative data. Yet, the consideration of time-point uncertainties substantially increases the computational complexity. The reasons are the increased model dimension, the numerical stiffness of the augmented model, in particular when time-point uncertainties are small, and the reduced convergence rate. These issues of the marginalized approach might for instance be addressed using more tailored numerical solvers. The efficiency of the joint approach could potentially be improved by using a hierarchical formulation of the estimation problem, similar to (Loos et al., 2018). However, the optimization for ***τ*** can be non-convex even if ***θ*** is known (see description of potential multimodality of the integrand in the *Methods* section).

The proposed approach relies on the assumption that the distribution of measurement time uncertainty is known or can be parameterized. In practice, this distribution may not be directly observable and may need to be inferred or approximated based on experimental protocols. While our framework allows for the estimation of such parameters (as showcased in the application problem), identifiability may depend on the amount and informativeness of the available data. Furthermore, the computational cost of evaluating the marginalized likelihood is increased compared to the standard formulation due to the required numerical integration. However, as demonstrated, this additional cost is offset by improved sampling efficiency and reduced dimensionality.

Our formulation is general and can be applied to a wide range of dynamic models and experimental settings. In particular, it is relevant for studies where measurement timing is uncertain due to experimental constraints or reporting inaccuracies, such as in clinical or high-throughput settings. In conclusion, accounting for measurement time uncertainty is essential for reliable parameter inference in dynamic models, and the marginalized formulation provides an efficient and scalable approach that combines statistical rigor with practical applicability.

## Author contributions

JV and JH designed the study. NB established the link to survival analysis. VN and JV implemented the models and performed the parameter estimation. JV and JH analyzed the data. JV and JH wrote the initial draft of the manuscript. All authors reviewed and approved the final version.

